# Role-Aware Line-Graph Neural Networks for Interpretable Prediction of Pairwise Microbial Interactions

**DOI:** 10.64898/2026.11.23.768606

**Authors:** Elham Gholamzadeh, Kajal Singla, Nico Scherf

## Abstract

Predicting how microbial species affect each other’s growth is a key challenge in microbial ecology. Using one of the largest available pairwise interaction datasets, comprising 20 soil bacteria co-cultured under 39 carbon conditions (15,074 directional measurements), we predicted whether one species has a positive or negative effect on the growth of another from monoculture growth, metabolic profiles, and phylogeny. Existing machine learning approaches treat each measurement as an independent sample, although experiments that share a species under the same condition are biologically related. We therefore represented the data as a role-aware line graph: each node is one directional measurement, and two nodes are connected when, under the same carbon condition, they share the same effector or the same affected species. This allows graph neural networks (GNNs) to pass information between related interactions. Our role-aware GAT-SAGE model achieved an F1-score of 81.38% for one-way prediction, outperforming the 73.79% previously reported for Extreme Gradient Boosting (XGBoost). For two-way interaction types (mutualism, competition, and parasitism), GAT-SAGE reached a macro F1-score of 60.34%. This exceeded the 55.71% of a neural network without graph structure but was below the 64.24% of XGBoost. Mutualism remained difficult to predict for all models. Layer-wise relevance propagation showed that one-way effects depend mainly on monoculture growth, whereas two-way interaction types depend more on the carbon environment. Encoding the dependency structure of interaction experiments thus adds information beyond the features of each measurement and provides interpretable insight into microbial interactions.

**IMPORTANCE:** Microbial communities in soil, in the human body, and in biotechnology are shaped by how their members affect each other’s growth, but measuring every pairwise interaction in every environment is impossible. Computational predictions can help to prioritize experiments, yet existing models treat each measured interaction as independent. We show that interaction measurements can instead be represented as a network in which related experiments are connected, so that graph neural networks can learn from neighboring measurements. This representation improves the prediction of interaction types compared with a neural network that ignores these relationships, and an interpretation of the trained models indicates that the growth of each species on its own determines the direction of an effect, whereas the environment shapes the type of reciprocal interaction. Positive and mutualistic interactions remain rare and hard to predict, which highlights where additional experimental data are most needed.

## INTRODUCTION

Understanding how microbial species function and interact is a fundamental problem in biology and biotechnology, and it remains experimentally challenging [22, 35]. A better understanding of these interactions would improve our ability to manage and engineer microbial communities, offering broad applications in environmental conservation, crop health, and human health [12, 33]. For several decades, various features of microbial communities have been exploited for the clean-up of contaminated water, soil, and air, and for the capture of renewable resources such as energy (biogas and H_2_), chemicals (short-chain acids), and water [31, 6]. The main advantages of microbial communities over pure cultures are non-sterile conditions, low-priced mixed or complex substrates, process robustness, and the ability to perform complex tasks based on metabolic modularity [26].

Interactions between species can be negative, where a species inhibits another species’ growth through nutrient exploitation and chemical warfare [7], or positive, where a species promotes another species’ growth by increasing nutrient availability and creating new niches [2]. In particular, positive, growth-promoting interactions can substantially affect the diversity and productivity of natural and engineered communities. Predicting interspecific interactions is therefore crucial for understanding a community’s properties, as they are shaped by the interactions within it. To understand and design interaction networks in larger communities, researchers often use co-culture experiments to measure pairwise interactions. However, because the number of possible pairwise combinations grows as the square of the number of species, and interactions also change with the environment, it is impossible to measure them exhaustively, even in moderately sized microbial communities.

Computational prediction of interactions could therefore greatly reduce the experimental effort. Predicting microbial interactions has generally depended on metabolic modeling and genome-based models [30, 16]. By taking into account the overlap and complementarity between species’ metabolic capacities and their resource consumption and secretion, these methods use genetic information to predict interactions [9, 14].

Machine learning offers a complementary, data-driven approach. Supervised machine learning algorithms [8, 29], including support vector machines [38], neural networks [28], random forests [4], and graph neural networks [24], have become popular for predicting complex phenotypes [3, 19, 11] due to the rise of high-throughput techniques that produce large biological data sets. In several biological domains, including microbiology, the use of supervised and unsupervised machine learning methods has grown in recent years [18, 27]. Prior research has demonstrated that deep learning can be used to predict the composition of microbial communities [1, 20]. Furthermore, based on genomic information and inferred metabolic pathways, it has recently been shown that supervised machine learning techniques can reliably predict the sign of microbial interactions (positive or negative) [5].

Building on the work of Nestor et al. [23], we leverage one of the largest data sets of experimentally validated microbial interactions to evaluate how well graph neural networks can predict the sign of microbial interactions (positive/negative) compared to other standard machine learning models. Similar to that work, we use species’ phylogeny, monoculture growth yield measured across each of the 40 carbon environments, and metabolic capacity as features to build our models. The data set comprises all pairwise interactions between 20 distinct soil bacteria from two taxonomic groups, cultivated in 40 different environments, each containing either a single carbon source or a combination of all carbon sources.

Existing machine learning approaches, however, typically treat these interaction measurements as independent samples. In reality, experiments that share a microbial species under the same environmental condition are biologically related. Moreover, interactions are directional: a species can act on others (as the effector) in some experiments and be affected by others in different experiments. A model that ignores this structure cannot use information from related experiments, and a model that mixes a species’ two roles may combine information that is biologically unrelated.

To address this limitation, we propose using graph neural networks (GNNs) as a data-driven model of microbial interactions, since GNNs can leverage information across separate but related experiments. We introduce a role-aware line-graph representation in which each node is a single directional interaction (effector *→* affected) under a specific carbon condition, and two nodes are connected only if they share a species playing the same role, either the same effector or the same affected species, under the same carbon condition. This enables message passing over interactions rather than species, while preserving the directionality of each interaction. We hypothesize that connecting measurements that share a species in the same role provides a useful inductive bias, allowing the model to learn how strongly a species tends to affect others, and how sensitive it is to others, from related experiments.

Figure **1** summarizes our framework. Co-culture experiments under each carbon condition (Figure **1**a) are described by features of the interacting species, including monoculture growth, metabolic profiles, and phylogeny (Figure **1**b). We first represent the experiments as a directed species graph, in which arrows denote the effect of one species on another (Figure **1**c). Each directional measurement is then turned into a node (Figure **1**d), and two nodes are connected only if they share the same effector or the same affected species under the same carbon condition, yielding the role-aware line graph *L*(*G*) (Figure **1**e). Finally, a GAT-SAGE model performs node classification on *L*(*G*), predicting either the sign of one-way interactions (binary classification) or the type of two-way interactions, i.e., mutualism, competition, or parasitism (multi-class classification) (Figure **1**f).

**FIG 1.**
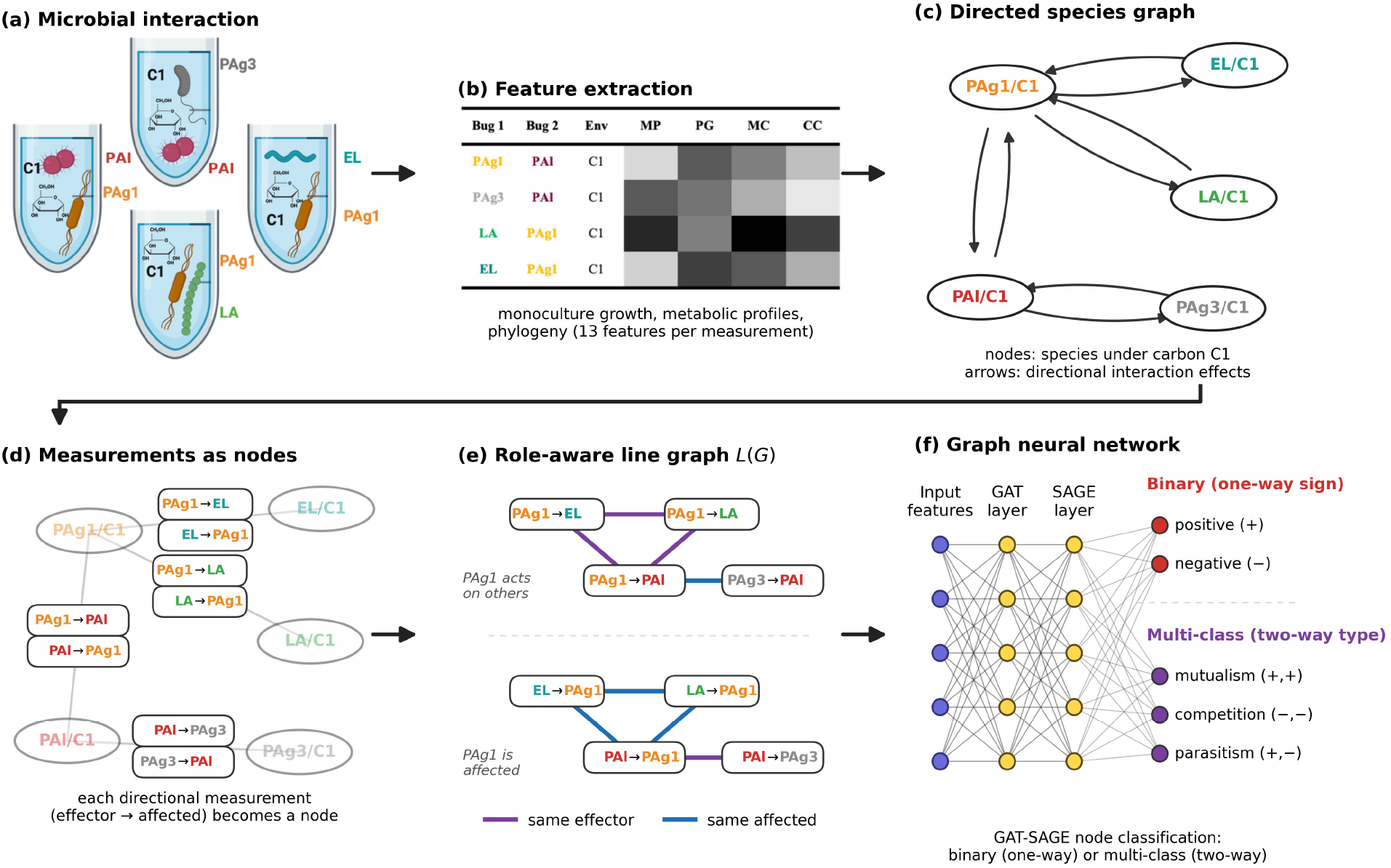
Overview of the role-aware line-graph framework for predicting microbial interactions. (a) Pairwise co-culture experiments under a carbon condition (C1). (b) Feature extraction for each measurement, including monoculture growth, metabolic profiles, and phylogeny (13 features per measurement). (c) Directed species graph: nodes are species under the same carbon condition, and arrows are directional interaction effects. (d) Each directional measurement (effector *→* affected) becomes a node. (e) Role-aware line graph *L*(*G*): two measurements are connected if they share the same effector (purple) or the same affected species (blue) under the same carbon condition. (f) The GAT-SAGE model (a graph attention layer followed by a GraphSAGE layer) classifies each node either as positive or negative (binary classification, one-way interaction) or as mutualism, competition, or parasitism (multi-class classification, two-way interaction).

Our main contributions are:

- a role-aware line-graph representation that connects directional interaction measurements sharing a species in the same role;
- a comparison of four GNN architectures (GraphSAGE, GAT, a hybrid GAT-SAGE model, and CGNN) for one-way and two-way interaction prediction, with an MLP without graph structure and XGBoost as baselines for two-way prediction, and an ablation of the role-aware graph;
- an analysis of class imbalance with random oversampling and entropy regularization; and
- an interpretation of the trained models with Layer-wise Relevance Propagation (LRP).

## RESULTS

### A role-aware line graph connects related interaction measurements

Each co-culture experiment yields two directional measurements, which we represented as the nodes of a line graph (Figure **1**; see Materials and Methods). In the role-aware line graph, two measurements are connected only if, under the same carbon condition, they share the same effector or the same affected species (Figure **7**). The resulting graph has 15,074 nodes and 277,719 edges, about half as many edges as a role-agnostic line graph that connects all measurements sharing any species (533,872 edges).

### GAT-SAGE performs best among the graph neural network architectures for one-way prediction

Among the four GNN architectures, the hybrid GAT-SAGE model achieved the highest F1-score for one-way sign prediction (81.38 *±* 2.21%), followed by GraphSAGE (79.70 *±* 2.77%), GAT (78.85 *±* 2.42%), and CGNN (76.38 *±* 2.33%) (Table **1**). GAT-SAGE also had the highest accuracy, recall, ROC-AUC, and PR-AUC. All four GNNs outperformed the 73.79% F1-score previously reported for Extreme Gradient Boosting (XGBoost) on the same task [23], with GAT-SAGE exceeding it by 7.6 percentage points. All GNNs classified negative measurements with few errors but missed some of the positive ones: GAT-SAGE detected 1,531 of the 1,960 positive measurements (78.1%) and produced 270 false positives (Figure **2**).

**TABLE 1.** Performance of role-aware GNN architectures, an MLP without graph structure and XGBoost for one-way interaction prediction (mean *±* standard deviation, *n* = 5 folds). The best value per column within each block is shown in bold. ^†^GAT-SAGE on the role-agnostic line graph, in which two measurements are connected if they share any species under the same carbon condition.

| Model | Accuracy | Precision | Recall | F1-score | ROC-AUC | PR-AUC |
|---|---|---|---|---|---|---|
| GAT-SAGE | **0.9533 ± 0.0070** | 0.8473 ± 0.0294 | **0.7830 ± 0.0166** | **0.8138 ± 0.0221** | **0.9694 ± 0.0039** | **0.8794 ± 0.0184** |
| GAT-SAGE<sup>†</sup> | 0.9514 ± 0.0059 | 0.8409 ± 0.0230 | 0.7730 ± 0.0223 | 0.8054 ± 0.0213 | 0.9672 ± 0.0035 | 0.8742 ± 0.0195 |
| GAT | 0.9477 ± 0.0069 | 0.8319 ± 0.0213 | 0.7495 ± 0.0270 | 0.7885 ± 0.0242 | 0.9588 ± 0.0026 | 0.8550 ± 0.0194 |
| GraphSAGE | 0.9502 ± 0.0073 | **0.8490 ± 0.0281** | 0.7516 ± 0.0341 | 0.7970 ± 0.0277 | 0.9638 ± 0.0051 | 0.8640 ± 0.0202 |
| CGNN | 0.9431 ± 0.0066 | 0.8300 ± 0.0235 | 0.7075 ± 0.0251 | 0.7638 ± 0.0233 | 0.9511 ± 0.0054 | 0.8236 ± 0.0189 |
| MLP | 0.8568 ± 0.0062 | 0.7750 ± 0.0171 | 0.5531 ± 0.0054 | 0.6454 ± 0.0072 | 0.8857 ± 0.0048 | 0.7636 ± 0.0107 |
| XGBoost | **0.8864 ± 0.0054** | **0.8089 ± 0.0123** | **0.6785 ± 0.0111** | **0.7379 ± 0.0086** | **0.9310 ± 0.0019** | **0.8447 ± 0.0080** |

**FIG 2.**
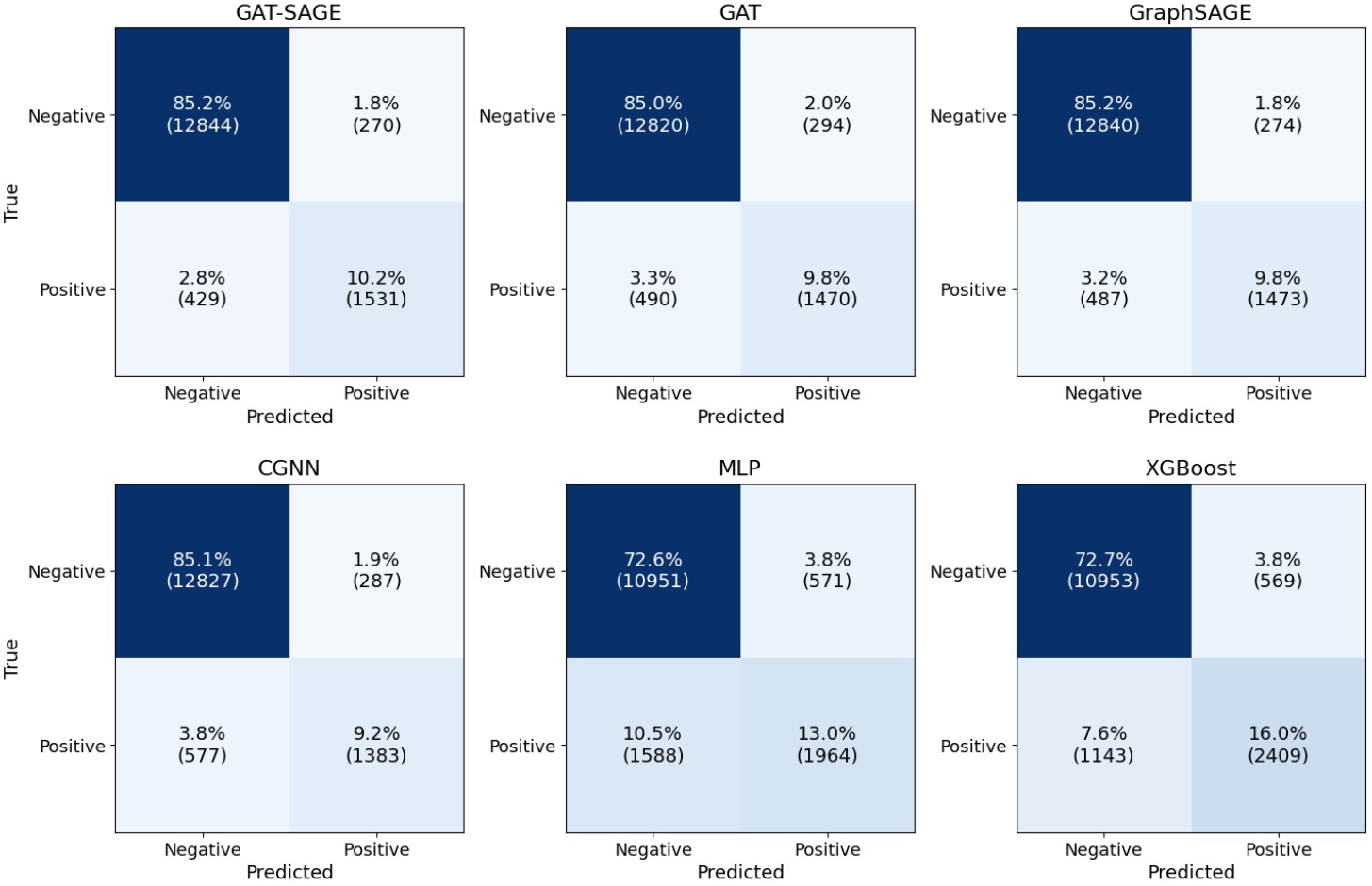
Confusion matrices for one-way sign prediction of the four GNN architectures, summed over the five validation folds. Each matrix is normalized by the total number of measurements, so that all cells sum to 100%; numbers in brackets are counts.

### Role-aware and role-agnostic line graphs perform similarly

For one-way prediction, GAT-SAGE reached an F1-score of 81.38 *±* 2.21% on the role-aware line graph and 80.54 *±* 2.13% on the role-agnostic line graph (Table **1**). The role-aware model also outperformed conventional Extreme Gradient Boosting (XGBoost) models, which reported an F1-score of 73.79%. Restricting edges to pairs that share a species in the same role therefore halved the number of edges without reducing performance.

### Message passing improves two-way prediction over a graph-free model, but XGBoost remains strongest

Table **2** and Figure **3** summarize the results for two-way interaction types. XGBoost achieved the highest macro F1-score (64.24 *±* 1.56%), followed by GAT-SAGE (60.34 *±* 1.10%), GraphSAGE (57.97 *±* 1.69%), the MLP (55.71 *±* 1.63%), GAT (54.51 *±* 1.80%), and CGNN (45.36 *±* 0.92%). Competition (*−*, *−*) was predicted well by all models; for GAT-SAGE, correctly classified competitive measurements accounted for 55.9% of all measurements (F1-score 0.87). Mutualism (+, +) was the most difficult class: GAT-SAGE correctly classified only 314 of 1,217 mutualistic measurements (2.2% of all measurements), and misclassified most of the remaining ones as parasitism (+, *−*) (765 measurements). XGBoost showed the same pattern, with a higher mutualism F1-score (0.41 versus 0.33). These errors are consistent with the difficulty of detecting positive one-way effects: a pair is mutualistic only if both directional effects are positive, so an error in either direction turns a mutualistic pair into a parasitic one.

**TABLE 2.** Classification performance for two-way microbial interactions (mean *±* standard deviation over 5 folds). Data from Nestor et al. [23]. XGBoost was trained with the hyperparameters of Nestor et al. [23], and the MLP is a neural network without graph structure; both use the same 13 features and folds as the GNNs. Best values per row are in bold. Prec.: precision; Rec.: recall.

| Class | Metric | GAT-SAGE | GAT | GraphSAGE | CGNN | MLP | XGBoost |
|---|---|---|---|---|---|---|---|
| Mutualism (+, +) | Prec. | 0.4607 ± 0.0575 | 0.4567 ± 0.0757 | 0.4262 ± 0.0576 | 0.3147 ± 0.1710 | 0.3728 ± 0.0625 | **0.4987 ± 0.0556** |
|  | Rec. | 0.2588 ± 0.0344 | 0.1203 ± 0.0450 | 0.1938 ± 0.0499 | 0.0264 ± 0.0148 | 0.1483 ± 0.0276 | **0.3473 ± 0.0457** |
|  | F1 | 0.3285 ± 0.0266 | 0.1859 ± 0.0614 | 0.2625 ± 0.0528 | 0.0484 ± 0.0269 | 0.2117 ± 0.0371 | **0.4065 ± 0.0384** |
| Competition (−, −) | Prec. | 0.8353 ± 0.0056 | 0.8147 ± 0.0130 | 0.8256 ± 0.0071 | 0.7399 ± 0.0062 | 0.8172 ± 0.0095 | **0.8553 ± 0.0068** |
|  | Rec. | 0.9095 ± 0.0125 | 0.9022 ± 0.0060 | **0.9155 ± 0.0112** | 0.8809 ± 0.0078 | 0.9121 ± 0.0090 | 0.9121 ± 0.0057 |
|  | F1 | 0.8708 ± 0.0057 | 0.8562 ± 0.0083 | 0.8682 ± 0.0075 | 0.8042 ± 0.0021 | 0.8621 ± 0.0083 | **0.8828 ± 0.0035** |
| Parasitism (+, −) | Prec. | 0.6310 ± 0.0158 | 0.5978 ± 0.0239 | 0.6314 ± 0.0129 | 0.5482 ± 0.0189 | 0.6197 ± 0.0106 | **0.6560 ± 0.0216** |
|  | Rec. | 0.5926 ± 0.0084 | 0.5890 ± 0.0298 | 0.5870 ± 0.0063 | 0.4737 ± 0.0022 | 0.5768 ± 0.0100 | **0.6220 ± 0.0171** |
|  | F1 | 0.6110 ± 0.0078 | 0.5931 ± 0.0244 | 0.6083 ± 0.0069 | 0.5081 ± 0.0083 | 0.5975 ± 0.0100 | **0.6380 ± 0.0072** |
| Macro F1 |  | 0.6034 ± 0.0110 | 0.5451 ± 0.0180 | 0.5797 ± 0.0169 | 0.4536 ± 0.0092 | 0.5571 ± 0.0163 | **0.6424 ± 0.0156** |
| Accuracy |  | 0.7594 ± 0.0054 | 0.7423 ± 0.0109 | 0.7559 ± 0.0059 | 0.6864 ± 0.0047 | 0.7471 ± 0.0095 | **0.7771 ± 0.0055** |

**FIG 3.**
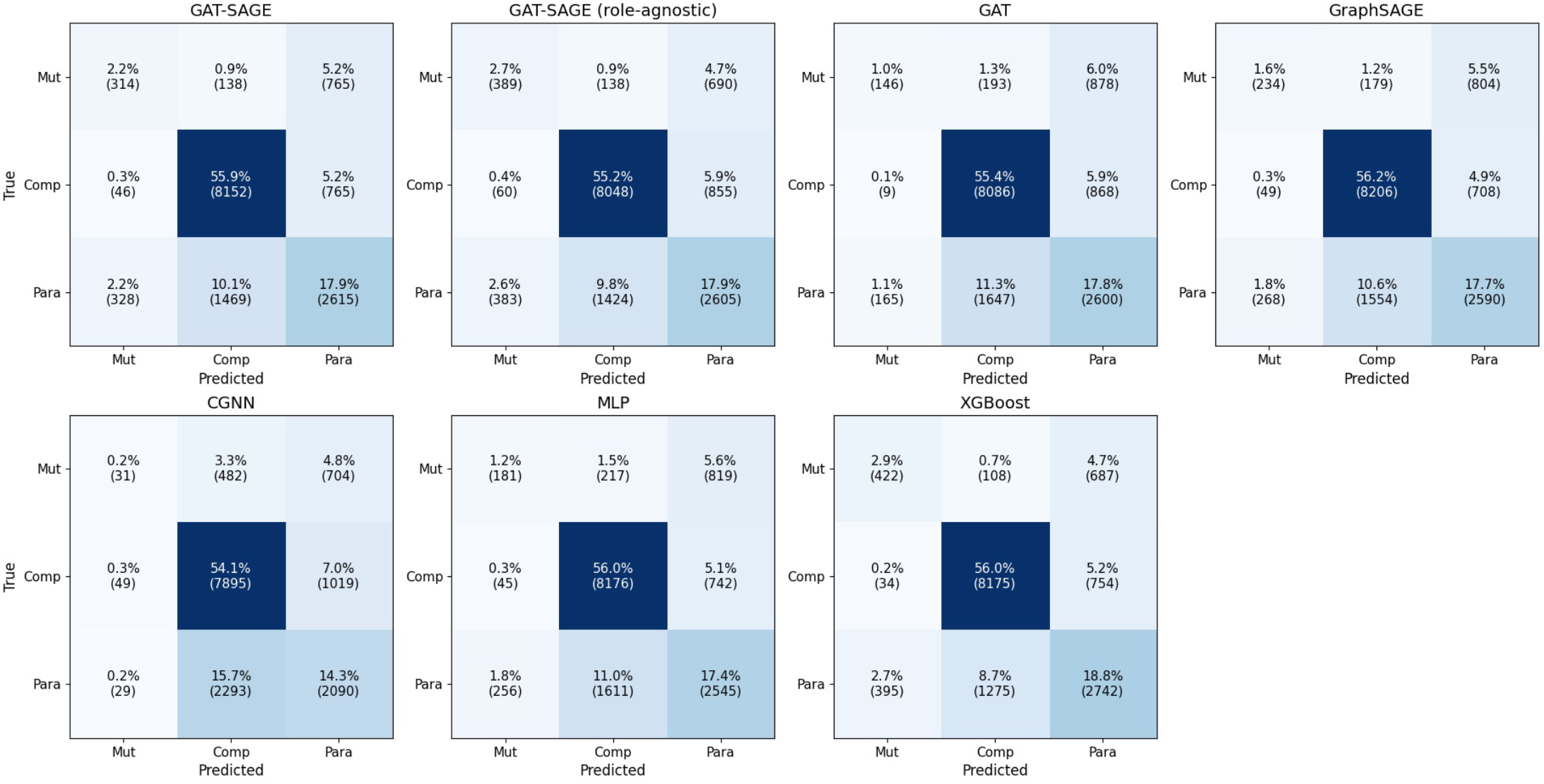
Confusion matrices for two-way interaction types (Mut: mutualism, Comp: competition, Para: parasitism) of the GNN architectures, GAT-SAGE on the role-agnostic line graph, the MLP, and XGBoost (hyperparameters of Nestor et al. [23]), summed over the five validation folds. Each matrix is normalized by the total number of measurements, so that all cells sum to 100%; numbers in brackets are counts.

### Oversampling trades precision for recall, whereas entropy regularization has little effect

Positive interactions are the minority class in one-way prediction, and all GNNs missed a part of them (Figure **2**). We therefore tested random oversampling and mixed entropy minimization with GAT-SAGE (**Tables 3** and **4**, Figure **4**).

**TABLE 3.** GAT-SAGE with mixed entropy minimisation for different balancing factors *λ* and with random oversampling of the minority class for one-way interaction prediction (mean *±* standard deviation, *n* = 5 folds; *λ* = 0 is plain cross-entropy). The best value in each column is shown in bold.

| *λ* | Accuracy | Precision | Recall | F1-score | ROC-AUC | PR-AUC |
|---|---|---|---|---|---|---|
| 0.0 | 0.9535 ± 0.0017 | 0.8483 ± 0.0167 | 0.7821 ± 0.0069 | 0.8138 ± 0.0095 | 0.9678 ± 0.0032 | 0.8770 ± 0.0103 |
| 0.1 | 0.9538 ± 0.0028 | 0.8497 ± 0.0217 | 0.7838 ± 0.0100 | 0.8153 ± 0.0115 | 0.9665 ± 0.0028 | 0.8773 ± 0.0088 |
| 0.2 | 0.9527 ± 0.0020 | 0.8450 ± 0.0221 | 0.7798 ± 0.0082 | 0.8108 ± 0.0078 | 0.9683 ± 0.0033 | 0.8802 ± 0.0031 |
| 0.3 | 0.9533 ± 0.0013 | 0.8529 ± 0.0233 | 0.7752 ± 0.0133 | 0.8118 ± 0.0058 | **0.9686** ± **0.0028** | 0.8790 ± 0.0059 |
| 0.4 | 0.9525 ± 0.0020 | 0.8443 ± 0.0259 | 0.7793 ± 0.0108 | 0.8101 ± 0.0071 | 0.9673 ± 0.0030 | 0.8769 ± 0.0077 |
| 0.5 | 0.9526 ± 0.0024 | 0.8497 ± 0.0270 | 0.7726 ± 0.0103 | 0.8090 ± 0.0096 | 0.9678 ± 0.0027 | **0.8828** ± **0.0075** |
| 0.6 | 0.9524 ± 0.0027 | 0.8470 ± 0.0287 | 0.7741 ± 0.0070 | 0.8086 ± 0.0115 | 0.9671 ± 0.0026 | 0.8792 ± 0.0119 |
| 0.7 | 0.9534 ± 0.0014 | 0.8567 ± 0.0187 | 0.7705 ± 0.0086 | 0.8111 ± 0.0064 | 0.9665 ± 0.0032 | 0.8791 ± 0.0098 |
| 0.8 | 0.9524 ± 0.0033 | 0.8511 ± 0.0274 | 0.7688 ± 0.0085 | 0.8077 ± 0.0153 | 0.9682 ± 0.0026 | 0.8803 ± 0.0103 |
| 0.9 | **0.9546** ± **0.0024** | **0.8635** ± **0.0236** | 0.7735 ± 0.0057 | **0.8158** ± **0.0109** | 0.9678 ± 0.0039 | 0.8804 ± 0.0134 |
| Oversampling | 0.9240 ± 0.0045 | 0.6527 ± 0.0212 | **0.8892** ± **0.0122** | 0.7526 ± 0.0161 | 0.9676 ± 0.0037 | 0.8694 ± 0.0069 |

**TABLE 4.**
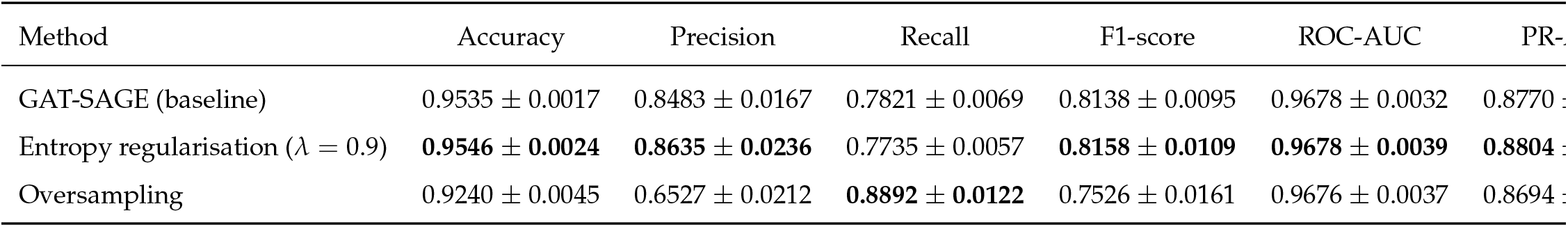
Strategies against class imbalance for GAT-SAGE, one-way interaction prediction (mean *±* standard deviation, *n* = 5 folds). The best value in each column is shown in bold.

**FIG 4.**
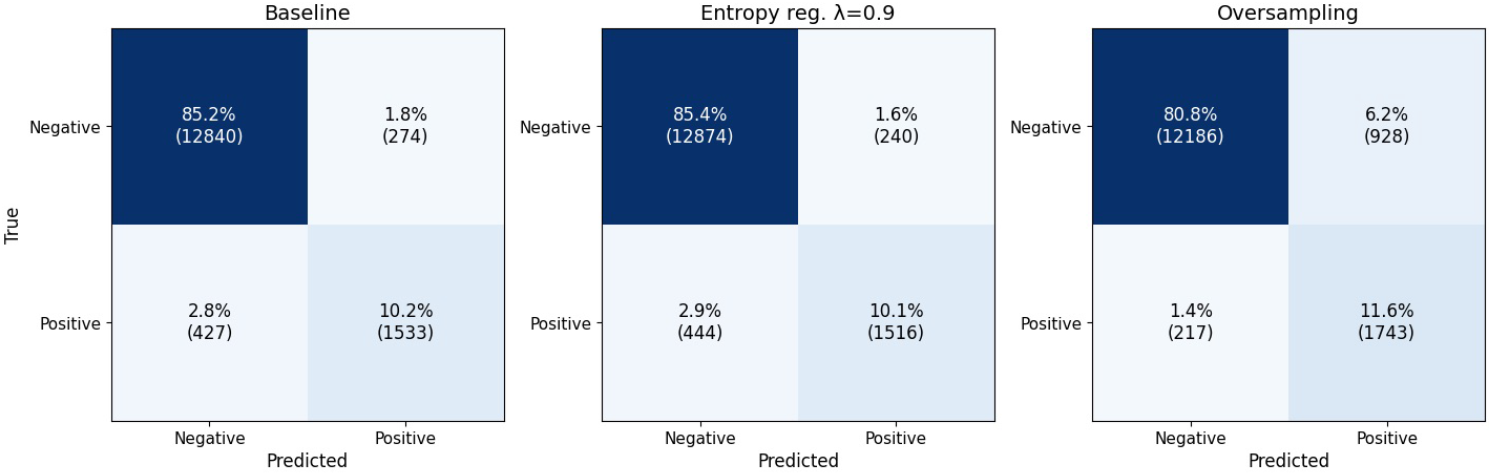
Confusion matrices of GAT-SAGE for one-way prediction without imbalance handling (baseline), with entropy regularization (*λ* = 0.9), and with random oversampling, summed over the five validation folds and normalized by the total number of measurements.

Entropy regularization had little effect. Across all balancing factors, the F1-score ranged from 80.77% to 81.58%, and the best setting (*λ* = 0.9, 81.58 *±* 1.09%) was within one standard deviation of the baseline without regularization (81.38 *±* 0.95%). Oversampling changed the balance between the two types of error: recall increased from 78.21 *±* 0.69% to 88.92 *±* 1.22%, and the number of correctly detected positive measurements rose from 1,533 to 1,743, but precision decreased from 84.83 *±* 1.67% to 65.27 *±* 2.12%, because the number of false positives increased from 274 to 928. As a result, the F1-score decreased to 75.26 *±* 1.61%. Oversampling is therefore useful when detecting as many positive interactions as possible matters more than avoiding false positives, for example when selecting candidate pairs for experimental validation.

### Different features drive one-way and two-way predictions

We used layer-wise relevance propagation (LRP) to estimate how much each input feature contributed to the predictions of GAT-SAGE, averaged over all nodes (Figures **5** and **6**).

**FIG 5.**
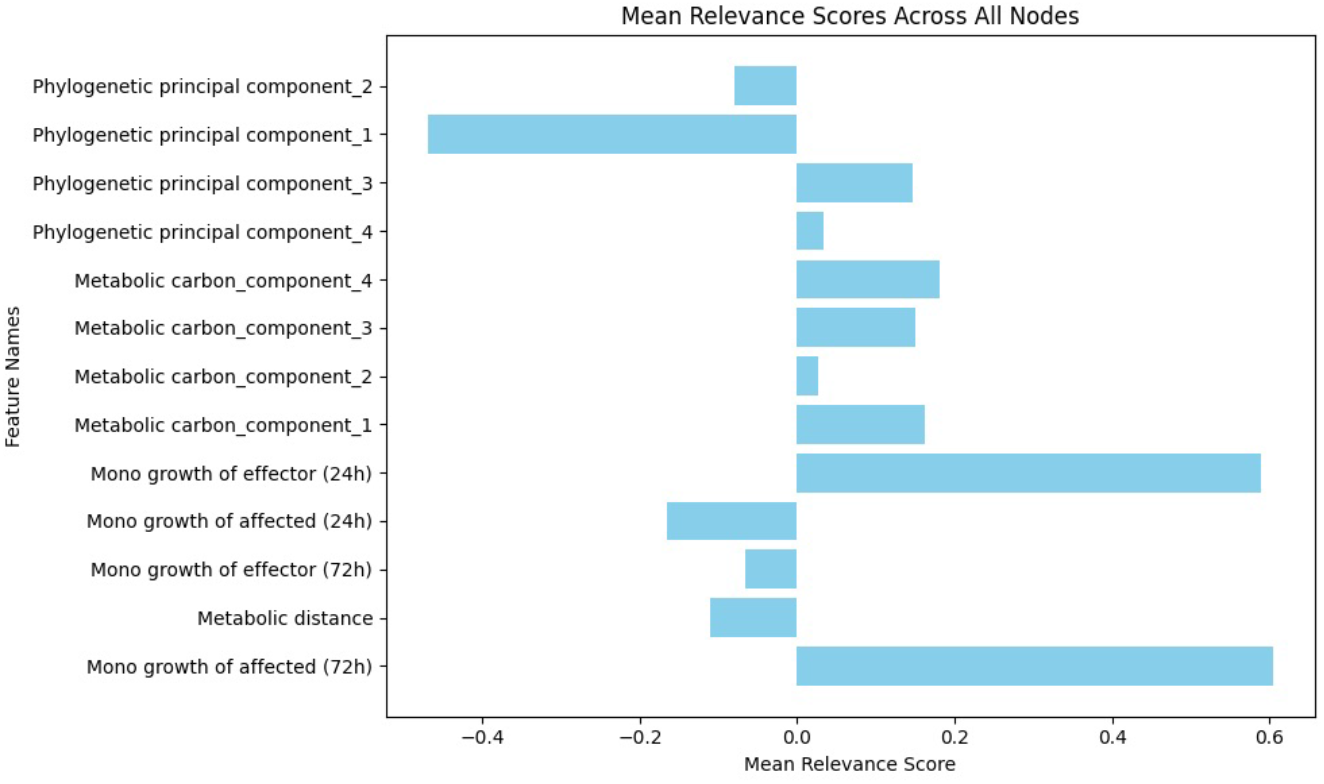
Mean relevance of each input feature for one-way sign prediction with GAT-SAGE, computed with layer-wise relevance propagation (LRP) and averaged over all nodes. Positive values indicate a contribution toward the predicted class and negative values a contribution against it. The monoculture growth of the affected species after 72 h and of the effector species after 24 h are the most relevant features.

**FIG 6.**
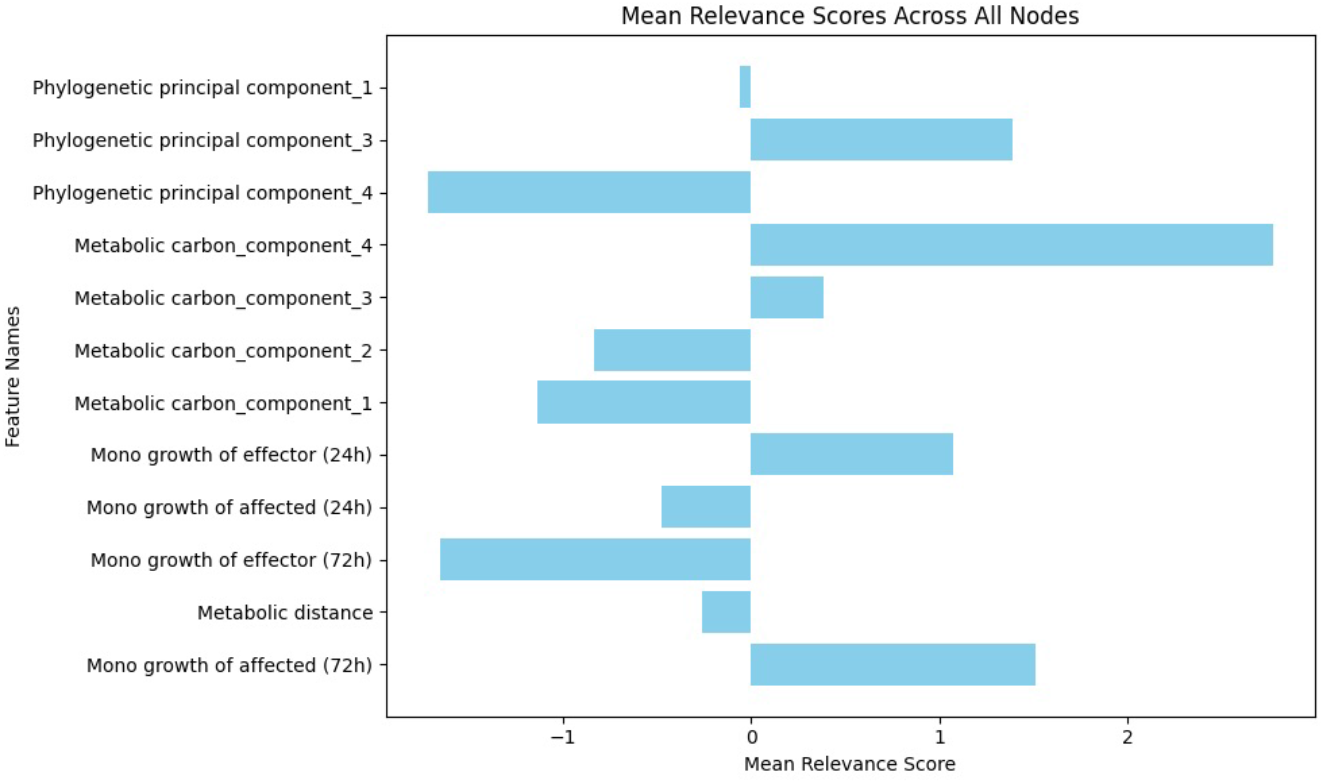
Mean relevance of each input feature for two-way interaction prediction with GAT-SAGE, computed with layer-wise relevance propagation (LRP) and averaged over all nodes. Positive values indicate a contribution toward the predicted class and negative values a contribution against it. The fourth principal component of the carbon-source metabolic profile and the monoculture growth of the affected species after 72 h are the most relevant features.

For one-way interaction prediction, the most relevant features were the monoculture growth of the affected species after 72 h and of the effector species after 24 h, whereas the first phylogenetic principal component contributed negatively to the model output (Figure **5**).

For two-way interaction prediction, the most relevant features were the fourth principal component of the metabolic profile of the carbon source and the monoculture growth of the affected species after 72 h (Figure **6**). Compared with one-way prediction, the relevance thus shifted from features of the individual species toward a feature that describes the carbon environment.

## DISCUSSION

### Encoding the structure of interaction experiments improves prediction

Most machine learning approaches for predicting microbial interactions treat each measurement as an independent sample [23, 5]. However, measurements in a combinatorial co-culture screen are not independent: every measurement shares its effector and its affected species, under the same carbon condition, with many other measurements. Our line-graph formulation makes this dependency explicit and allows GNNs to pass information between related measurements. The clearest test of this idea is the two-way task, in which all models were trained and evaluated on the same labels and folds: GAT-SAGE reached a macro F1-score of 60.34%, compared with 55.71% for an MLP with the same features and model capacity but without message passing. Because the two models differ only in the graph, this difference isolates the contribution of message passing and indicates that information from related experiments, and not only the features of the measurement itself, improves the neural model.

The role-aware construction was designed to refine this idea. In a role-agnostic line graph, the neighborhood of a measurement mixes experiments in which a shared species acts on others with experiments in which it is affected by others. In the role-aware line graph, these two roles are separated (Figure **7**): the neighbors with the same effector inform the model about how strongly that species tends to affect others, and the neighbors with the same affected species inform it about how sensitive that species is to others. Both are biologically meaningful properties of a species in a given environment. In practice, however, the role-aware and role-agnostic graphs performed similarly: for one-way prediction, the F1-score of GAT-SAGE was 81.38% with the role-aware and 80.54% with the role-agnostic graph, a difference smaller than one standard deviation across folds. This indicates that the models are robust to the exact definition of the graph. The role-aware graph is nevertheless about half as dense (277,719 versus 533,872 edges) and easier to interpret, because each edge has a clear biological meaning.

**FIG 7.**
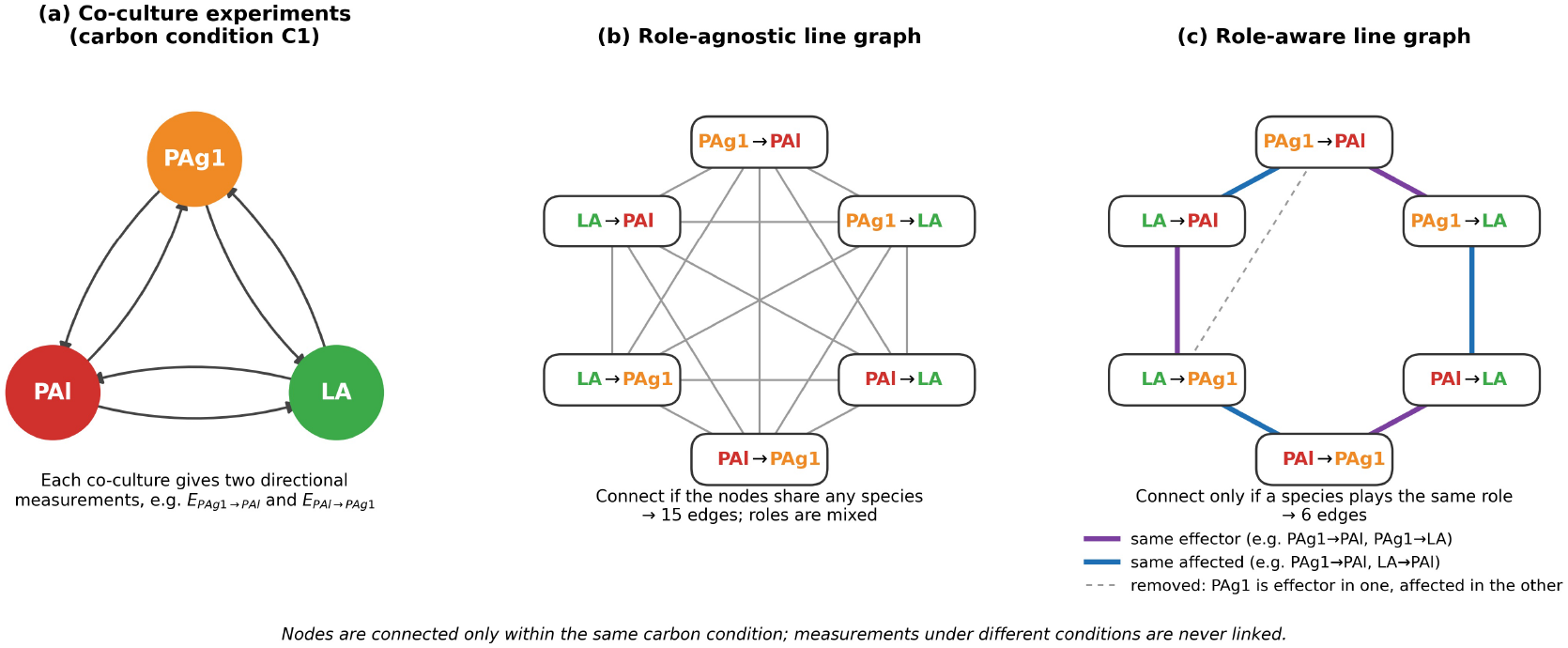
Role-aware line-graph construction, illustrated for three species (PAg1, PAl, LA) under carbon condition C1. (a) Each co-culture experiment yields two directional measurements, e.g., *E*_PAg1_*_→_*_PAl_ and *E*_PAl_*_→_*_PAg1_. (b) In the role-agnostic line graph, two measurements are connected whenever they share any species, mixing a species’ effector and affected roles. (c) In the role-aware line graph, two measurements are connected only if they share the same effector (purple) or the same affected species (blue); a species acting as effector in one measurement and as affected in the other does not create an edge (dashed, removed). Nodes are connected only within the same carbon condition.

Among the architectures, the hybrid GAT-SAGE model performed best for one-way prediction (Table **1**). Attention allows the model to weight the most informative related measurements, while mean aggregation in the subsequent SAGE layer provides a stable summary of the neighborhood. CGNN performed worst, suggesting that edge curvature, which reflects how strongly the neighborhoods of two measurements overlap, is less informative for this task than the features of the neighbors themselves. Because nodes in the role-aware line graph form dense groups of measurements that share a species under one condition, most edges have similar curvature, which limits the additional information that curvature-based weighting can provide.

### Two-way interactions remain difficult to predict

For two-way interaction types, XGBoost remained the strongest model, and competition was predicted well by all models, whereas mutualism was the most difficult class and was mostly misclassified as parasitism (Table **2**, Figure **3**). This is expected for two reasons. First, mutualism is rare in the data (8.3% of pairs), so the models see few examples of it. Second, a pair is mutualistic only if both directional effects are positive, so any error in either direction turns a mutualistic pair into a parasitic one. The difficulty of predicting mutualism therefore follows directly from the difficulty of predicting positive one-way effects, which are themselves the minority class. Neither oversampling nor entropy regularization resolved this imbalance: in one-way prediction, oversampling increased the recall of GAT-SAGE from 78.21% to 88.92% but reduced its precision from 84.83% to 65.27%, and entropy regularization changed the F1-score only marginally (81.58% at *λ* = 0.9 versus 81.38% without it). That positive and mutualistic interactions are rare and hard to predict is itself an ecologically relevant finding, consistent with the observation that competition dominates pairwise interactions between culturable bacteria [7].

### Species growth and the environment shape different aspects of interactions

Feature relevance analysis showed that different features drive the two prediction tasks (Figures **5** and **6**). For one-way prediction, the most relevant features were the monoculture growth of the affected species after 72 h and of the effector species after 24 h. This indicates that the effect of one species on another depends mainly on how well each species grows on its own in the given carbon condition. A plausible interpretation is that a species that grows well alone has little room to be promoted and is more likely to be inhibited by competition for the same resource, whereas a species that grows poorly alone can benefit more from metabolites released by a fast-growing partner. Consistent with this, Nestor et al. [23] also identified monoculture growth as one of the most informative features.

For two-way prediction, the most relevant features included a principal component of the metabolic profile of the carbon source, together with the monoculture growth of the affected species. Because this component describes the carbon environment rather than the species, this result suggests that the type of reciprocal interaction (mutualism, competition, or parasitism) depends strongly on the environment, for example on whether a carbon source can be used by both species, leading to competition, or is converted into by-products that one species can use, allowing cross-feeding. The shift from species-specific growth features in one-way prediction to environmental features in two-way prediction suggests that the direction of an effect is determined mainly by the traits of the two species, whereas the overall type of the interaction emerges from how both species use the available resources.

### Limitations and outlook

Our study has several limitations. First, all interactions were measured in vitro under controlled laboratory conditions, and it remains unclear how well the predictions transfer to natural environments with fluctuating conditions and spatial structure. Second, the models predict pairwise interactions and do not capture higher-order effects, in which the interaction between two species is modified by a third. Third, the line-graph formulation relies on information about related measurements and on monoculture growth of each species in each condition; predictions for species or conditions without such data are therefore less reliable, and generalization to entirely unseen species remains to be tested. Fourth, GNNs are computationally more demanding than tree-based models such as XGBoost, which remained the strongest baseline in our comparisons. Finally, our evaluation is transductive: the full graph is available during training, which reflects the practical use case of predicting missing measurements in a partially measured screen, but not the prediction of interactions for completely new communities.

Several extensions could address these limitations. The role-aware line graph could be generalized to a heterogeneous graph with separate edge types for shared effectors, shared affected species, and opposite roles, which a relational GNN could weight differently. Higher-order interactions could be modeled with topological deep learning approaches that represent groups of more than two species. Finally, applying the framework to other interaction datasets, and to other systems with directional links such as host–pathogen or epidemiological networks, would test its generality.

### Conclusions

In this work, we showed that an edge-focused graph learning approach can capture both the structure and the direction of interactions in a complex biological system. By converting a combinatorial co-culture screen into a role-aware line graph, in which each node is a directional interaction measurement and edges connect measurements that share a species in the same role under the same carbon condition, graph neural networks learn the properties of interactions directly, rather than only the traits of individual species. Information from related experiments improved prediction compared with a neural network without graph structure, and the hybrid GAT-SAGE model performed best among the GNN architectures, although XGBoost remained a strong baseline for two-way interaction types.

Layer-wise Relevance Propagation revealed that monoculture growth plays a central role in predicting the direction of an effect, whereas the carbon environment contributes more to the type of reciprocal interaction. Oversampling and mixed entropy minimization did not fully resolve the class imbalance, and positive and mutualistic interactions remained underrepresented and difficult to predict. Notably, this aligns with the ecological observation that such interactions are genuinely rare among culturable bacteria [7].

The method is generalizable and could be applied to other ecological systems, epidemiological networks, or any network with diverse, directional links. In the long run, by embedding biologically meaningful features at each edge, geometric deep learning models can achieve high predictive accuracy while reflecting real-world interaction patterns, bridging the gap between data-driven prediction and biological understanding.

## MATERIALS AND METHODS

### Dataset

We used the pairwise interaction dataset of Nestor et al. [23], which is based on the kChip platform of Kehe et al. [13], a high-throughput, nanodroplet-based system for combinatorial screening of species growth in monoculture and co-culture. The dataset comprises all pairwise co-cultures of 20 soil bacteria from two taxonomic groups across 40 environments: single carbon sources (some at two concentrations), a mixture of all carbon sources, and a water control. We excluded the water control, leaving 39 carbon conditions. Each co-culture of species *A* and *B* yields two directional measurements: the effect of *B* on the growth of *A*, *E_B→A_*, and the effect of *A* on the growth of *B*, *E_A→B_*. In total, the dataset contains 15,074 directional interaction measurements.

### Features

Following Nestor et al. [23], each measurement is described by 13 features (Table A.1 in Appendix): the monoculture growth yield of both species after 24 and 72 hours in the given carbon condition, the metabolic distance between the two species, the first four principal components of the metabolic profile of the carbon source (describing the environment), and the first two principal components of the phylogenetic distance matrix for each species. All features were min–max scaled.

### Interaction labels

#### One-way interactions

Each directional measurement is labeled positive if its effect *E_B→A_* is greater than zero and negative otherwise, resulting in 23.6% positive and 76.4% negative measurements.

#### Two-way interactions

For each species pair *A* and *B* under a carbon condition, the signs of both directional effects are combined,

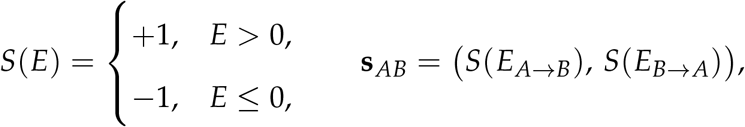

and the interaction type is defined as mutualism if **s***_AB_* = (+1, +1), competition if **s***_AB_* = (*−*1, *−*1), and parasitism otherwise. Because the interaction type is a property of the pair, both directional measurements of a pair carry the same two-way label, resulting in 8.3% mutualism, 61.4% competition, and 30.2% parasitism.

### Role-aware line-graph construction

#### Species graph

For each carbon condition, we define a directed species graph *G* = (*V*, *E*). Each node *v ∈ V* is a species–carbon combination, *v* = (species, carbon) (e.g., PAg1/C1), and each directed edge (*B*, *A*) *∈ E* represents the measured effect of species *B* (the effector) on species *A* (the affected species) in co-culture (Figure **1**c).

#### Line graph

Our prediction targets are the interactions, i.e., the edges of *G*. We therefore transform *G* into its line graph *L*(*G*) = (*V^′^*, *E^′^*), in which every directional measurement becomes a node (Figure **1**d):

- **Node mapping:** each directed edge (*B_k_ → A_k_*) under carbon condition *C_k_* becomes a node 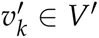.
- **Feature mapping:** the 13 features of the measurement are assigned to 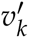.
- **Adjacency mapping (role-aware):** two nodes 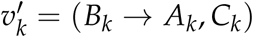 and 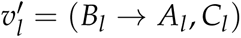 are connected if and only if

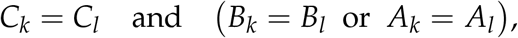

that is, the two measurements were taken under the same carbon condition and share a species that plays the *same* role in both: the same effector or the same affected species.

A species that acts as effector in one measurement and as affected species in another does not create an edge. In this way, message passing combines information from related experiments while keeping a species’ effect on others separate from the effect of others on it (Figure **7**). The resulting line graph contains 15,074 nodes and 277,719 edges.

#### Role-agnostic line graph (ablation)

To test the effect of the role-aware rule, we also built a role-agnostic line graph in which two nodes are connected whenever they share any species under the same carbon condition, regardless of its role (Figure **7**b). This graph contains 533,872 edges, almost twice as many as the role-aware graph.

### Graph neural network models

GNNs are message passing neural networks [36]: in each layer, the representation of node *i* is updated from its own features **x***_i_* and an aggregation of the features of its neighbors *N*(*i*),

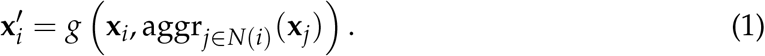

We compared four two-layer architectures that differ in how neighbors are aggregated. All models perform node classification on *L*(*G*), binary for one-way and three-class for two-way interactions, and are trained with the cross-entropy loss

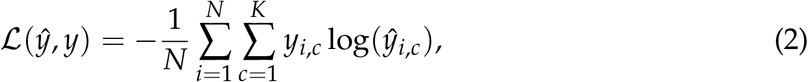

where *N* is the number of training nodes, *K* is the number of classes (*K* = 2 or *K* = 3), *y_i_*_,*c*_ is the one-hot true label, and *ŷ_i,c_* is the predicted probability.

#### GraphSAGE

GraphSAGE [10] combines a node’s own features with the mean of its neighbors’ features,

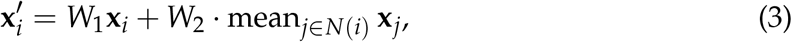

where *W*_1_ and *W*_2_ are learnable weight matrices.

#### Graph attention network (GAT)

GAT [32] weights each neighbor by a learned attention coefficient,

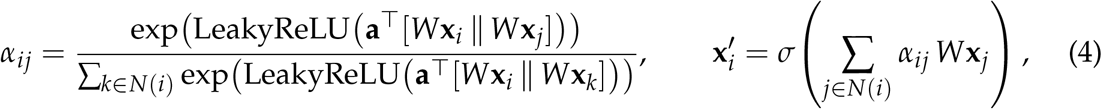

where **a** is a learnable attention vector and *σ* is a non-linear activation.

#### GAT-SAGE

To combine adaptive neighbor weighting with robust mean aggregation, we built a sequential hybrid model consisting of a multi-head GAT layer followed by a GraphSAGE layer,

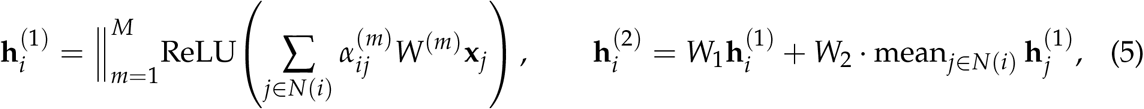

where the outputs of the *M* = 4 attention heads are concatenated (||). The attention layer first weights related interactions, and the SAGE layer then aggregates these representations into the final node prediction (Figure **1**f).

#### Curvature graph neural network (CGNN)

CGNN [17] weights messages by the Ollivier–Ricci curvature *r_ij_* of each edge, symmetrically normalized as 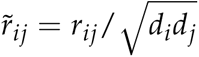 with 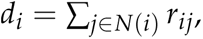

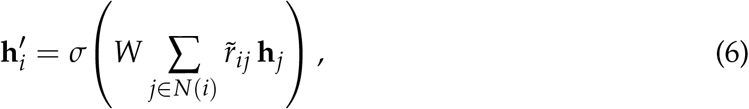

with dropout between the two layers. Curvature reflects how strongly the neighborhoods of two connected nodes overlap, so CGNN adapts message passing to the local geometry of the graph.

#### Baselines and ablations

To test whether the graph structure contributes to two-way prediction, we compared the GNNs with reference models trained on the same 13 features and evaluated on the same folds: (i) Extreme Gradient Boosting (XGBoost), the best-performing model of Nestor et al. [23]; and (ii) a multilayer perceptron (MLP) with the same depth and hidden size as the GNNs but without message passing, which isolates the contribution of the graph. In addition, we compared the role-aware with the role-agnostic line graph using GAT-SAGE.

### Class imbalance

Positive one-way interactions and mutualistic two-way interactions are underrepresented. We tested two strategies to address this imbalance. First, we randomly oversampled the minority class: in each fold, training nodes of the smaller class were drawn again with replacement until both classes had the same number of training nodes, while the graph and the validation nodes were left unchanged. Second, we used mixed entropy minimization (ME) [37], which adds the predictive entropy of the training predictions as a regularization term to the cross-entropy loss,

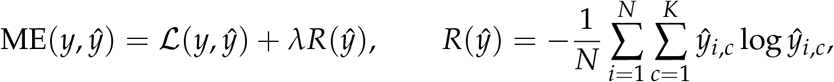

where the balancing factor *λ* was varied from 0 to 0.9.

### Training and evaluation

All models were implemented in Python using the Deep Graph Library (DGL) [34] and PyTorch [25]. The GNNs and the MLP used a hidden dimension of 100 and no dropout, and were trained with the Adam optimizer [15] with a learning rate of 10*^−^*^3^ for 3,00 epochs.

We used 5-fold cross-validation with a fixed random seed in a transductive setting: the full graph *L*(*G*) and all node features are available during training, and only the labels are split into training and validation folds. To prevent the two directional measurements of the same co-culture from appearing in different folds, folds were grouped by co-culture (unordered species pair and carbon condition) and stratified by class using StratifiedGroupKFold from scikit-learn. All models, including the baselines, were evaluated on the same folds.

We report the mean and standard deviation over the five folds of accuracy, precision, recall, and F1-score, and for one-way prediction also the areas under the ROC and precision–recall curves (ROC-AUC and PR-AUC; definitions in Appendix). Because the classes are imbalanced, we use the F1-score of the positive (minority) class as the main metric for one-way prediction, and the per-class and macro F1-scores for two-way prediction. To inspect the learned representations, we projected the hidden-layer embeddings of the validation nodes into two dimensions using t-SNE, for both one-way and two-way prediction. All experiments were run on the RAVEN HPC system (Intel Xeon IceLake-SP processors, NVIDIA A100 GPUs).

### Feature relevance

To interpret the predictions of the GAT-SAGE model, we used a relevance propagation procedure based on Layer-wise Relevance Propagation (LRP) [21]. Starting from the output activation of a target node, relevance is propagated backward through the graph layers: at each layer, it is redistributed through the layer weights to the node itself and, via message passing, to its neighbors, with a small constant *ɛ* added for numerical stability and a normalization step to prevent exploding values. This yields a relevance score for each input feature of each node. We report the mean relevance of each feature over all nodes, separately for the one-way and the two-way model.

## Data availability

The interaction data analyzed in this study were published by Nestor et al. [23] and are available at https://github.com/einatnestor/Microbial-interaction-prediction. The code for the line-graph construction, the graph neural network models, and the class-imbalance experiments is available at https://github.com/elhamgholamzadeh/role-aware-line-graph-microbial-interactions.

## ACKNOWLEDGMENTS

Computations were performed on the RAVEN HPC system of the Max Planck Computing and Data Facility.

## FUNDING

Our research is supported by the European Union and the Free State of Saxony through BIOWIN.

## AUTHOR CONTRIBUTIONS

E.G. and N.S. conceived the study. E.G., K.S., and N.S. developed the methodology. E.G. implemented the software, performed the analyses, prepared the visualizations, and wrote the manuscript. N.S. supervised the project, acquired funding, and reviewed and edited the manuscript. All authors read and approved the final manuscript.

## CONFLICTS OF INTEREST

The authors declare no conflicts of interest.

## SUPPLEMENTAL MATERIAL

### Features names information

Table A.1 summarizes the features used for microbial interaction prediction, including monoculture growth measurements, metabolic dissimilarity, and principal components derived from carbon source utilization and phylogenetic embeddings.

**TABLE A.1.**
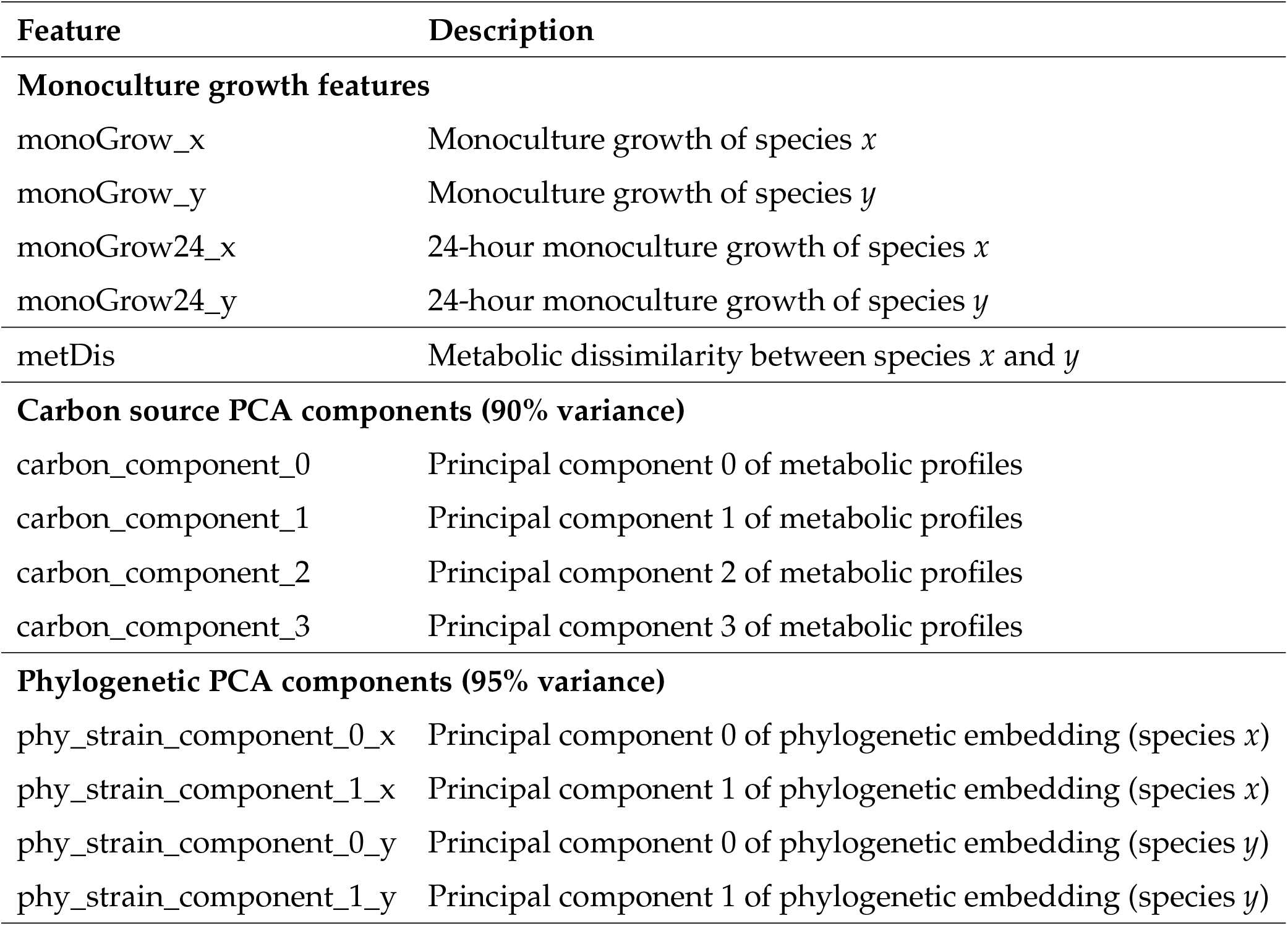
Features used for microbial interaction prediction.

| **Feature** | **Description** |
|---|---|
| **Monoculture growth features** |  |
| monoGrow_x | Monoculture growth of species *x* |
| monoGrow_y | Monoculture growth of species *y* |
| monoGrow24_x | 24-hour monoculture growth of species *x* |
| monoGrow24_y | 24-hour monoculture growth of species *y* |
| metDis | Metabolic dissimilarity between species *x* and *y* |
| **Carbon source PCA components (90% variance)** |  |
| carbon_component_0 | Principal component 0 of metabolic profiles |
| carbon_component_1 | Principal component 1 of metabolic profiles |
| carbon_component_2 | Principal component 2 of metabolic profiles |
| carbon_component_3 | Principal component 3 of metabolic profiles |
| **Phylogenetic PCA components (95% variance)** |  |
| phy_strain_component_0_x | Principal component 0 of phylogenetic embedding (species *x*) |
| phy_strain_component_1_x | Principal component 1 of phylogenetic embedding (species *x*) |
| phy_strain_component_0_y | Principal component 0 of phylogenetic embedding (species *y*) |
| phy_strain_component_1_y | Principal component 1 of phylogenetic embedding (species *y*) |

### Evaluation metric definitions

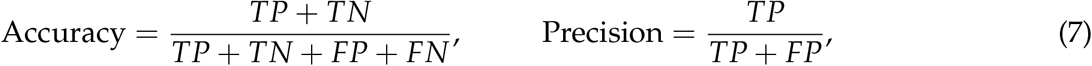

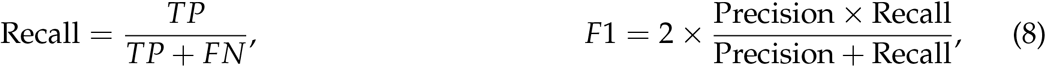

where TP, TN, FP, and FN are the numbers of true positives, true negatives, false positives, and false negatives. For two-way prediction, these metrics are computed for each class against all others.

Figures 8 and **9** show t-SNE projections of the hidden-layer embeddings of the validation nodes, for one-way and two-way prediction. For XGBoost, which has no hidden layer, the 13 input features are shown. In all models, the projections consist of several separate groups of measurements, and the classes are mixed within these groups rather than forming separate clusters. In the one-way embeddings of the GNNs, positive measurements tend to form small local groups within the larger clusters; in the two-way embeddings, mutualistic and parasitic measurements are interspersed with competitive ones. The two-dimensional projections therefore do not separate the classes clearly, and differences in prediction performance between the models (**Tables 1** and **2**) are not apparent from them. Because t-SNE preserves local rather than global structure, these plots should be interpreted qualitatively.

**FIG 8.**
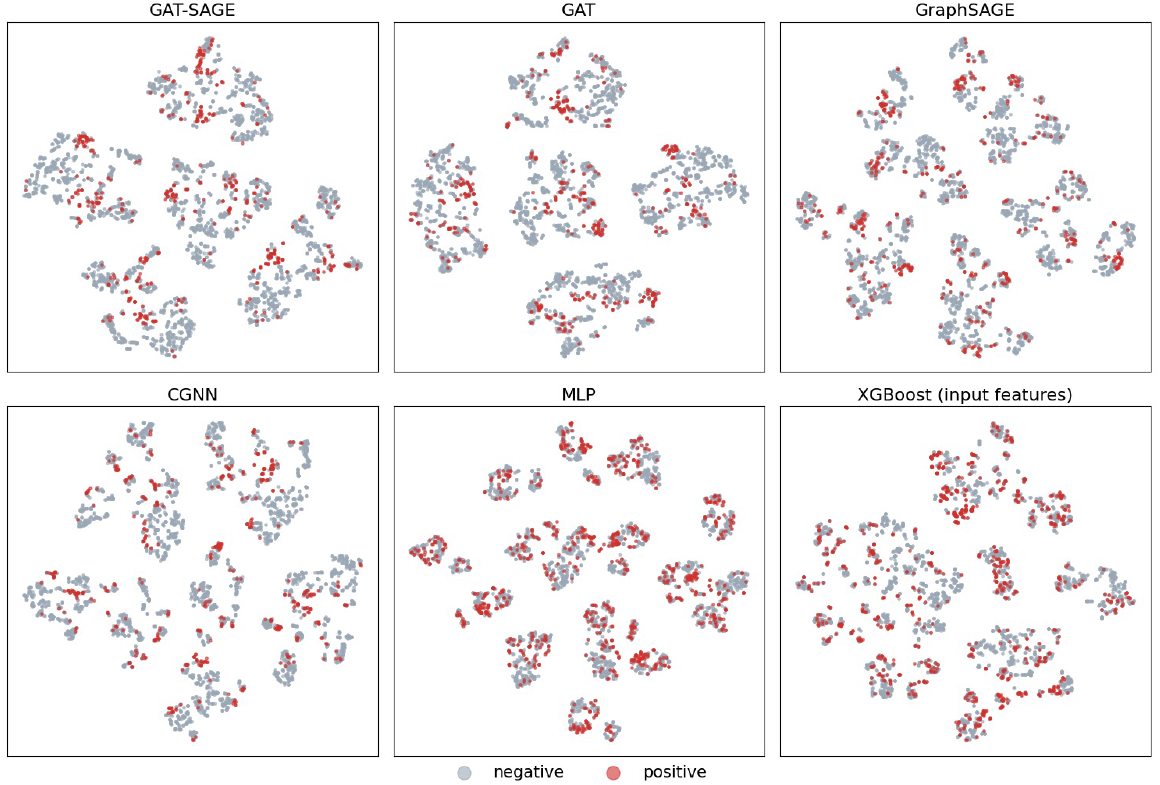
t-SNE projection of the hidden-layer embeddings of the validation nodes for one-way prediction (GAT-SAGE, GAT, GraphSAGE, CGNN, and MLP; 4,000 randomly sampled nodes). For XGBoost, the 13 input features are shown. Colors denote negative and positive interactions.

**FIG 9.**
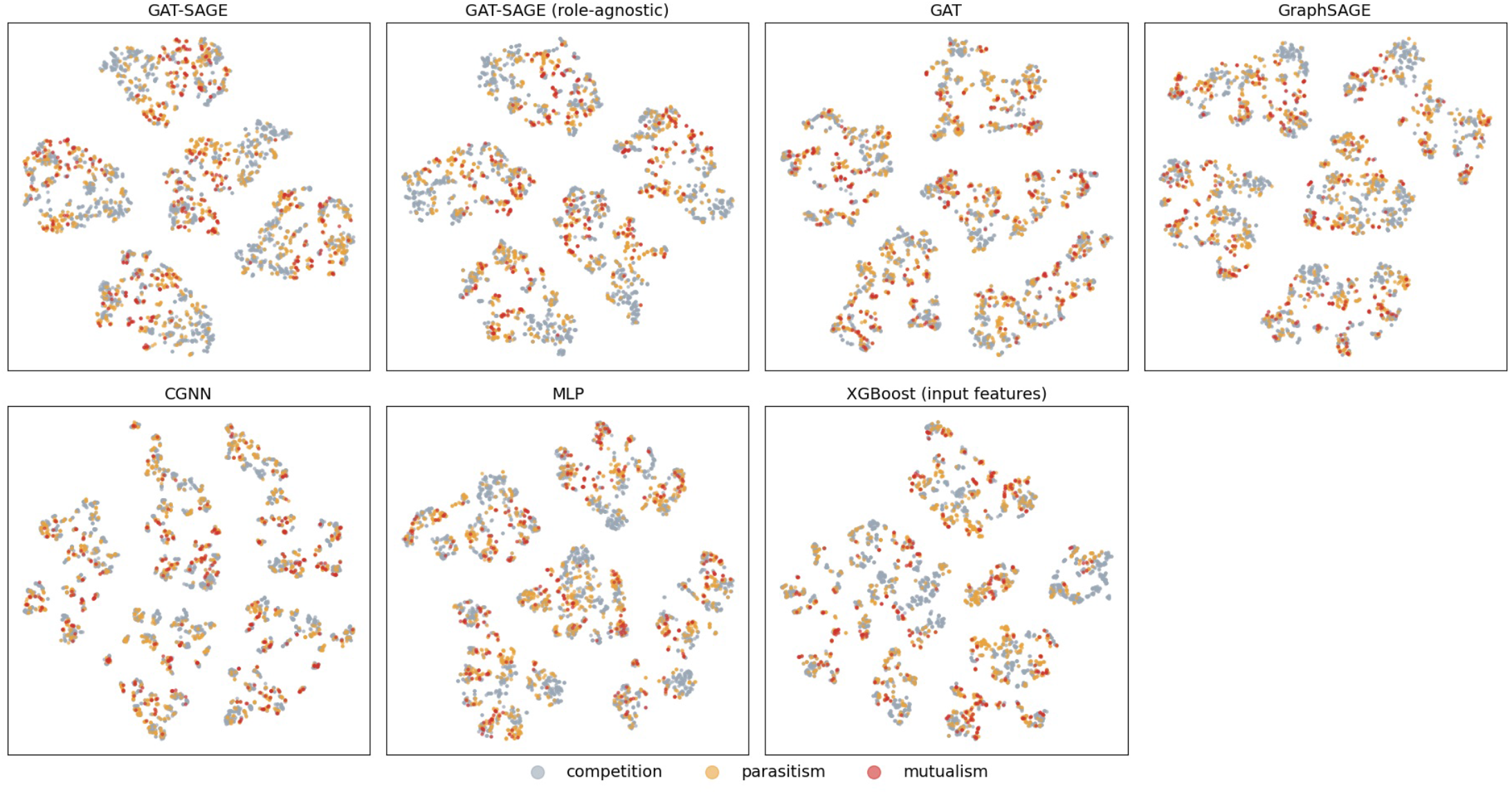
t-SNE projection of the hidden-layer embeddings of the validation nodes for two-way prediction (GAT-SAGE on the role-aware and role-agnostic line graphs, GAT, GraphSAGE, CGNN, and MLP; 4,000 randomly sampled nodes). For XGBoost, the 13 input features are shown. Colors denote competition, parasitism, and mutualism.

